# Species identity rather than colony size partitions shape variation in six Caribbean octocorals

**DOI:** 10.64898/2026.11.02.763628

**Authors:** Samuel Rivera-Muñiz, Christopher D. Wells, Howard R. Lasker

## Abstract

Caribbean reefs are increasingly dominated by arborescent octocorals as scleractinian cover declines. The physical structure of these octocoral forests is established by the manner in which octocoral species generate and modify colony form as they grow. We tested whether colony shape exhibited species-specific architecture and whether it changed allometrically with size. Two-dimensional projections of colony form were modeled as a series of stacked bands, whose areas were measured from scaled *in situ* images of colonies of six sympatric Caribbean octocorals from St John, US Virgin Islands. Principal component analysis (PCA) of relative band areas resolved a single dominant axis, accounting for 67.9% of the variation, that separated top-heavy from bottom- heavy colony profiles. Species identity explained 42% of the total variation in relative form, whereas colony height only explained 2% and had no detectable effect on either PCA axis. The proportion of the colony outline occupied by branches increased with height, from a predicted 41% at 20 cm to 52% at 100 cm. Colonies therefore grew without a detectable redistribution of area along their vertical axis, filling a species-specific outline more densely rather than moving along a common allometric trajectory. The vertical geometry of an octocoral canopy may therefore depend more on which species are present than on the sizes its colonies have reached.

## Introduction

Foundation species build the architecture of terrestrial and marine ecosystems, and many foundation species, including trees, macroalgae, sponges, bryozoans, hydroids, and corals, are modular, built from the iterative addition of structurally equivalent units (Harper and White 1974; Harper 1977). Modules may be the leaves and branches of vascular plants, the tillers and fronds of clonal grasses and algae, or the polyps, zooids, and whole branches of colonial invertebrates (Kaandorp and Kübler 2001; Lasker et al. 2003; Sánchez 2004). Because module addition continues through much of an individual’s life, modular organisms typically display indeterminate or loosely determinate growth and can reach sizes and ages that far exceed those of unitary organisms (Brazeau and Lasker 1988; Edmunds 2006; Shaish et al. 2006). Morphology in a modular organism is therefore the accumulated product of module generation and growth. It is this process, its astogeny (*sensu* Ruedemann 1904), not a pre-specified adult form, that any account of modular form must explain.

Modular organisms routinely build the three-dimensional matrix of the ecosystems in which they live. The same functional role is filled by different modular taxa in different systems: on land by trees in forest canopies, in temperate marine systems by kelp and other large macroalgae, and on tropical reefs by sponges and corals, where the resulting assemblages are often described as animal forests (Rossi et al. 2017; Orejas et al. 2022). General principles linking the architecture of arborescent bodies to their ecological function are well developed in plant ecology through allometric and hydraulic scaling theory (King 1990; Enquist 2002) and through biophysical models of how tropisms and elasticity shape canopy outlines (Duchemin et al. 2018; Moulia et al. 2022). An analogous framework for colonial marine animals is considerably less developed, despite the architectural parallels long noted between marine animal forests and their botanical counterparts (Rossi et al. 2017; Orejas et al. 2022).

In this report, we focus on arborescent octocoral forests on Caribbean reefs. Scleractinian cover has declined steeply across the Caribbean (Jackson et al. 2014; Tebbett et al. 2023), while the abundance of arborescent octocorals has remained stable or increased on many reefs (Ruzicka et al. 2013; Lenz et al. 2015; Sánchez et al. 2019; Lasker et al. 2020, 2026). The resulting assemblages have been described as octocoral forests, three-dimensional canopies that provide habitat for reef organisms (Tsounis et al. 2020), alter near-bed flow (Guizien and Ghisalberti 2017) and sedimentation (Cerpovicz and Lasker 2021), and restructure the understory benthic community in ways broadly analogous to the effects of stony-coral frameworks (Girard and Edmunds 2023). On these reefs the dominant modular architects are shifting in relative abundance, but the ecological role of modular architecture as the habitat-building element has not.

Translating the shift in dominance into a prediction about reef function requires understanding how form varies among species and how that form develops. Modular colonies grow by the iterative processes of branching, polyp origination, and branch elongation, so the canopy a colony presents to the water column at any moment is the cumulative product of a developmental program rather than a fixed adult body plan (Lasker and Sánchez 2002; Sánchez 2004). The modification of conditions within and around a colony and the habitat it provides are properties of that program, and they change as the colony grows. Colonies that look alike at one size could diverge at another, and a single colony may fill different ecological roles at different points along its growth trajectory. Establishing how colony form develops and how species-specific rules of module deployment generate the canopy at successive sizes is therefore a prerequisite to mapping the morphometrics of present-day octocorals onto the ecological role of the assemblages they build. The same logic underlies the developmental and allometric framing of plant canopy ecology (King 1990; Enquist 2002; Barthélémy and Caraglio 2007) and applies equally to colonial animals.

Octocorals are a particularly tractable system in which to study colony astogeny. Colonies develop from a single founding polyp and grow through the iterated origination of polyps, their organization into branches, and the formation and extension of branches, producing a discrete, free-standing arborescent body that can be measured in situ from scaled photographs (Lasker et al. 2003; Goffredo and Lasker 2006). Multiple closely related species co- occur across the same Caribbean reefs and span a wide range of mature architectures and sizes, ranging from open, sparsely branched whips to densely branched plumate forms, to net-like fans created by the fusion of branches (Bayer 1961; Kupfner Johnson and Hallock 2020). Colony morphology in these taxa is genetically structured but not rigid, and substantial within-species variation is generated by plastic responses to flow, light, depth, and physical disturbance (Kim et al. 2004; Prada and Hellberg 2013). Competitive interactions with canopy neighbors (Gambrel and Lasker 2016) together with the accumulated effects of partial mortality and regeneration (Castanaro and Lasker 2003; Sánchez and Lasker 2004a; Anderson et al. 2022) add further variance to colony form. Crucially, because colonies grow through module addition, form is inseparable from size, and any colony provides a snapshot along an astogenetic trajectory that characterizes the colony’s growth (Lasker and Sánchez 2002; Sánchez 2004; Shaish et al. 2006). In long-lived sessile taxa, following the trajectory of a single colony would require decades of study, but because colonies of different sizes represent different developmental stages, cross-sectional morphometrics, treating size as a proxy for astogenetic time (Wenker and Stevens 2020), as is standard in plant allometry (King 1990; Enquist 2002), allow the growth trajectory of form to be inferred from images of colonies spanning a range of sizes. Taken together, these features make Caribbean octocorals a natural bridge between the well-developed architectural frameworks of plant ecology and the less-developed ones of colonial animals.

Here, we use quantitative measurements of colony form, extracted from scaled *in situ* images of 52 colonies distributed across six Caribbean octocoral species, to study modular morphogenesis. We test two hypotheses framed around form generation in modular organisms: (1) species differ in relative colony morphology independent of size, such that the rules governing branch origination and placement produce species-specific architectures that remain recognizable at any developmental stage; and (2) relative colony form changes systematically with colony height, indicating that module deployment is allometric and that larger colonies occupy a different region of shape space than smaller conspecifics. Partitioning colony morphology in this way separates a species-specific component from a shared, continuous consequence of growth, and illustrates how cross-sectional morphometrics applied to an arborescent cnidarian can inform broader questions about how form develops in modular organisms.

## Methods

*In situ* photographs of 52 octocoral colonies from six species; *Pseudoplexaura wagenaari*, *Pseudoplexaura tenuis* (formerly *Pseudoplexaura crucis*), *Plexaura homomalla*, *Eunicea flexuosa* (formerly *Plexaura flexuosa*), *Antillogorgia americana* (formerly *Pseudopterogorgia americana*), and a currently undescribed species in the genus *Antillogorgia* similar in form but distinct from *Antillogorgia americana* (hereafter *Antillogorgia* sp. 1) were obtained at five fringing reefs on the southern shore of St John, US Virgin Islands. The sites lie within a single shallow fore-reef system, and colonies were photographed during the boreal summers of 2021, 2022, and 2024. Because colony form is the product of many years of growth, colonies were pooled across the three seasons. Site was included as a factor in the analyses of relative form and of branch fraction. Six to fifteen colonies were measured per species, and they ranged from 12 to 112 cm tall. We photographed only free-standing colonies that were not interacting with neighbors and showed no signs of previous partial mortality, so that measured form reflected undisturbed growth. Species were identified in the field, and photographs were taken by positioning a Plexiglas board with a 10 cm × 10 cm grid behind colonies so images could be scaled and measured (Fig. 1A; Online Fig. 1A). Distortion created when the camera was not perpendicular to the plane of the colony profile was corrected using Adobe Photoshop (v. 21.2), as in Lasker et al. (2003). Briefly, a digital grid was overlaid on the image, and the image’s shape was adjusted using the free transform function to align the 10-cm grid in the photograph with the digital grid. After images were aligned and scaled, Photoshop’s polygonal lasso tool was used to create polygons describing the perimeter of each colony by starting at the base and tracing the edge of the colony until the first branch tip was reached, then a convex hull of the branches was created to standardize the tracing of complex colony forms (Fig. 1B). With the use of the measurement tool, colony height (cm), width (cm), and area (cm²) were obtained for each colony. The polygon of each colony was then divided into ten stacked horizontal bands of equal height based on the total height of the colony (Fig. 1C) with band 1 starting at the colony’s base and band 10 ending at the top of the colony. The area and width of the portion of the polygon falling within that band was then determined using the measurement tool. Each colony image was then uploaded to CoralNet (Beijbom et al. 2015), which was used to determine the proportion of the polygon area occupied by branches. For each colony, 250 randomly distributed points were classified as either: on a colony branch, inside the colony polygon but not on a branch, or outside the polygon (Fig. 1D). Between 105 and 184 points fell inside the polygon of a given colony, and the branch fraction of that colony is the number of branch points out of that total.

**Fig. 1.**
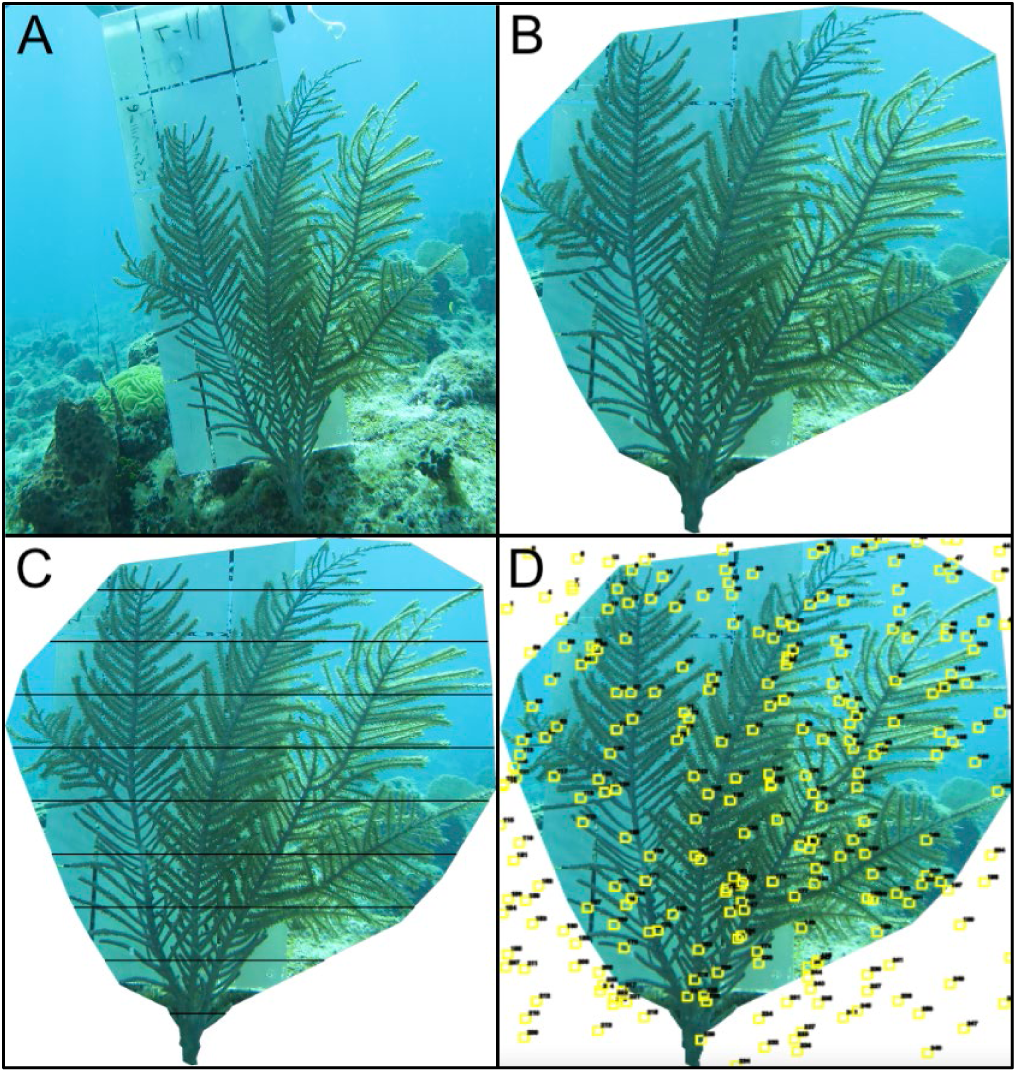
Steps involved in the analysis of images and collection of morphometric data for a colony of *Antillogorgia americana*. A, Initial image with a 10 × 10 cm scaling grid; B, Colony polygon; C, Colony bands; D, CoralNet point counts.

All statistical analyses were performed in R v4.6.0 (R Core Team 2026). Relative band areas were obtained by dividing each band area by the sum of the ten band areas of that colony, so that the ten relative areas of a colony sum to one. Principal component analyses (PCA), with each band standardized to unit variance, were conducted on relative band areas, to summarize variation in colony form independent of overall size, and separately on absolute band areas, to evaluate how colonies’ use of space varied across colony height and species.

Differences in species shape were tested using permutational multivariate analysis of variance (PERMANOVA; R package vegan 2.7-3, Oksanen et al. 2026) on relative band areas with Euclidean distances and 9,999 permutations, treating species identity as a fixed factor. Every permutation test reported here used the random- number seed 20260908. Homogeneity of multivariate dispersion among species was evaluated using a permutational analysis of multivariate dispersions (PERMDISP). PC1 and PC2 scores were tested separately using one-way linear models (PC ∼ Species), with significance assessed using analysis of variance (ANOVA) and the two p-values adjusted with a Bonferroni correction. Pairwise differences between species were tested via PERMANOVA on relative band areas and significance was adjusted across the 15 pairs by the Benjamini- Hochberg false-discovery-rate (FDR) procedure. Site was tested as a factor in a further PERMANOVA on relative band areas that also included species and colony height, with each term tested after the others. Because the ten relative band areas of a colony are a closed composition, the species and height PERMANOVAs were repeated on centered log-ratio transformed band areas.

To test whether relative colony form changed with colony size, ordinary least squares linear regressions were fit between colony height (cm) and each of the first two principal component scores derived from the relative band area PCA (PC1 ∼ Height; PC2 ∼ Height). To evaluate whether size-shape relationships differed among species, additional models including species identity and its interaction with height were fit (PC ∼ Height × Species), and significance of main effects and interactions was assessed using ANOVA. Slopes of PC1 on height for each species were estimated as marginal trends from the interaction model (R package emmeans 2.0.3, Lenth and Piaskowski 2026). A complementary multivariate test of height effects on relative morphology was performed using PERMANOVA on relative band areas (relative band areas ∼ Height) with Euclidean distances and 9,999 permutations, and repeated with species in the model (relative band areas ∼ Species + Height), testing each term after the other. The proportion of polygon area occupied by branches was modeled from the CoralNet point counts using a generalized linear model with a logit link and quasi-binomial errors. The quasi-binomial variance accommodated the overdispersion that arises because points within a colony fall on the same branches. The reported model has site as a factor and colony height as a covariate. A second model added species and a height- by-species interaction to test whether branch fraction increased differently among species.

## Results

Principal component analyses of absolute and relative band areas provided two views of morphological variation (Fig. 2). For absolute band areas, small colonies had similar morphologies regardless of species. PC1 explained 88.3% of the variance, and as all bands made similar contributions to the PC1 scores, the axis primarily reflected colony size. In combination, the first two axes accounted for 96.8% of total variation with the additional variance being explained by differences in the factor loadings of the different band heights, i.e., colony shape. In contrast, PCA of relative band areas summarized variation in colony form independent of size, with PC1 and PC2 explaining 67.9% and 18.7% of the variance, respectively (86.6% cumulative). Species separated along PC1 in the PCA based on relative band area and overlapped along PC2, indicating differences in the proportional distribution of colony area among height bands.

**Fig. 2.**
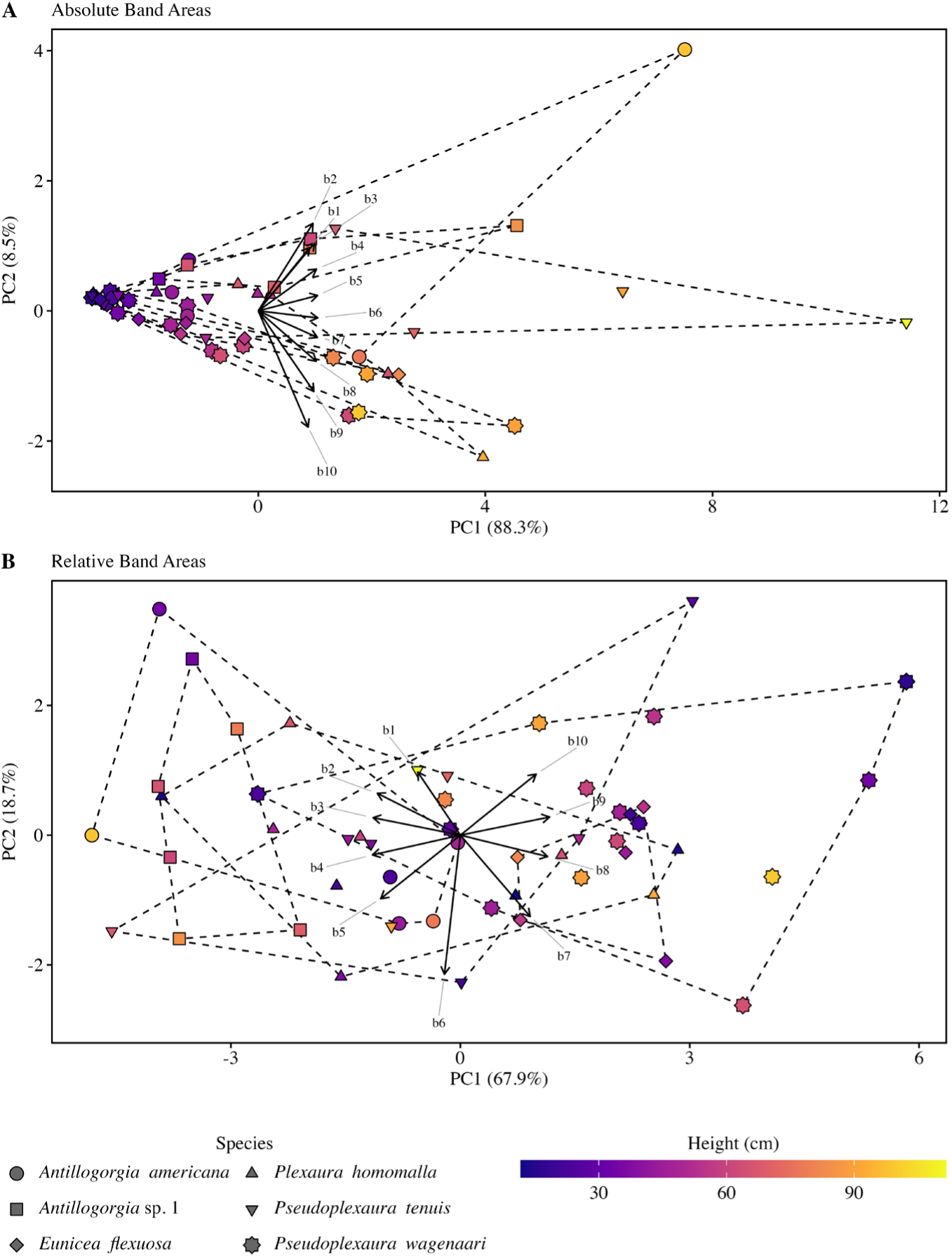
PC1 and PC2 scores from principal component analyses of absolute band areas (A) and relative band areas (B). Vectors illustrate the factor loadings for the ten bands.

A test for homogeneity of multivariate dispersions detected no difference among species in morphological variability (PERMDISP: *F*_5,46_ = 1.05, *p* = 0.40), consistent with the PERMANOVA assumption. Species differed significantly in relative band area profiles (PERMANOVA: *R*^2^ = 0.42, pseudo-*F*_5,46_ = 6.74, *p* = 0.0001), reflecting differences in their form independent of colony size (Fig. 3). Species were strongly differentiated along PC1 (67.9% of variance; *F*_5,46_ = 8.85, Bonferroni-adjusted *p* < 0.001) but did not differ along PC2 (18.7% of variance; *F*_5,46_ = 0.42, Bonferroni-adjusted *p* = 1.00). Relative factor loadings (vectors in Fig. 2B) showed that PC1 contrasted lower bands (b1-b5, with negative loadings) against upper bands (b7-b10, with positive loadings), so position along PC1 reflected whether colony area was concentrated relatively high (positive PC1, top-heavy) or extended into the lower bands (negative PC1, bottom-heavy). Pairwise PERMANOVA with false-discovery-rate correction indicated that eight of 15 species pairs were distinguishable in relative form (*p*_FDR_ ≤ 0.038). Every pair contrasting one of the two *Antillogorgia* species with either *E. flexuosa* or *P. wagenaari* was distinguishable, as were *Antillogorgia* sp. 1 from *P. homomalla* and *P. tenuis*, and *P. wagenaari* from those same two species. On PC1, *P. wagenaari* (mean = +1.98 ± 0.57 SE; all species means reported as mean ± SE) and *E. flexuosa* (+1.83 ± 0.35) occupied the top-heavy extreme, while *A. americana* (−1.81 ± 0.83) and *Antillogorgia* sp. 1 (−3.32 ± 0.29) occupied the bottom-heavy extreme, with *P. homomalla* (−0.56 ± 0.72) and *P. tenuis* (−0.47 ± 0.70) intermediate.

**Fig. 3.**
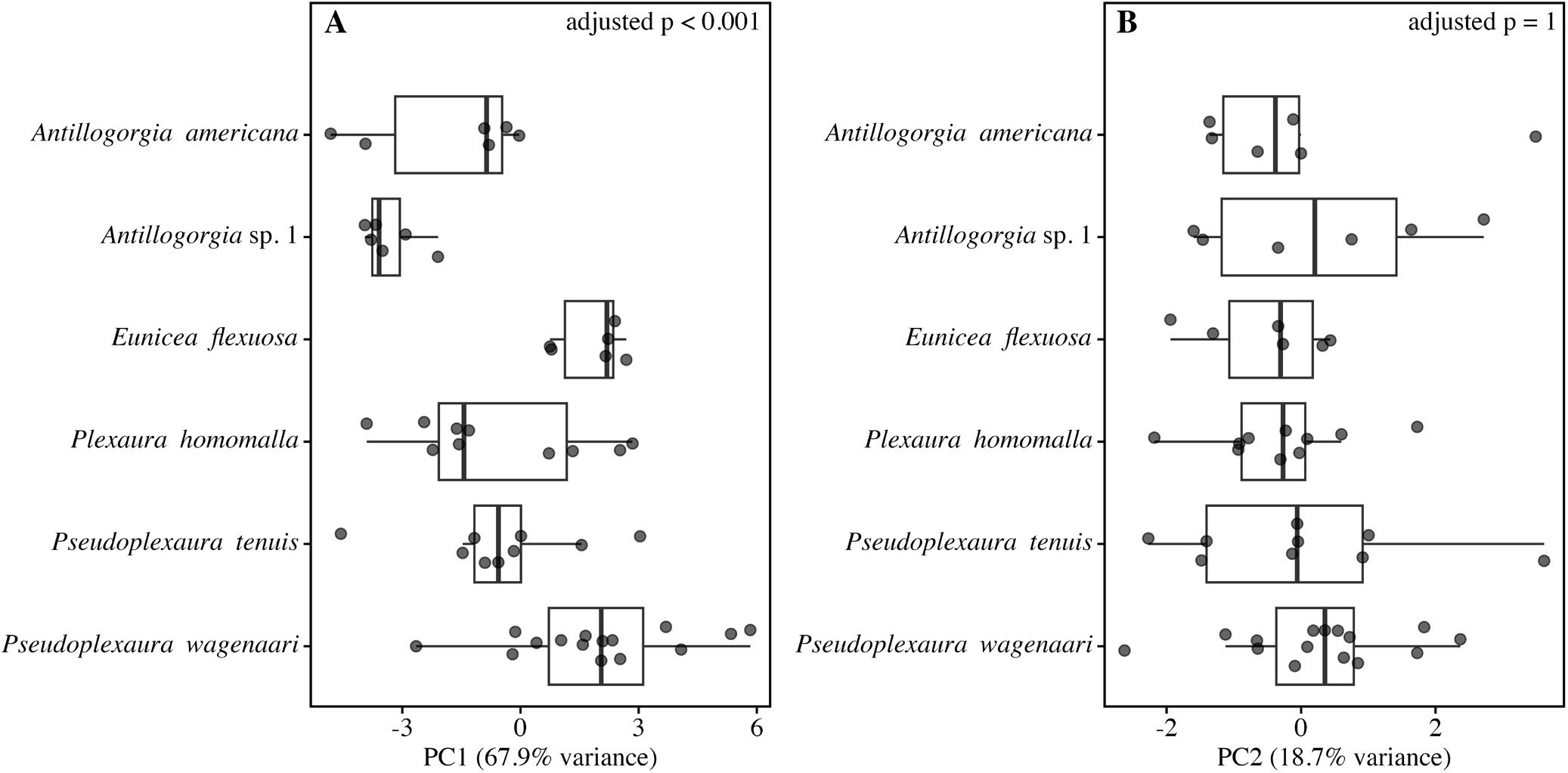
Box plots of species scores on the first two principal components using relative areas. Species separate on PC1 but overlap on PC2.

Average colony profiles for the six species were made using the average of each relative band area for every individual of the given species (Online Fig. 2). The profiles of all six species had parabola-like forms. *E. flexuosa and P. wagenaari* had nearly identical, top-heavy profiles, with 45.7 ± 1.0% and 45.9 ± 1.5% of relative colony area in bands 7-9 (60 to 90% of colony height); the two *Antillogorgia* species shared a more bottom-heavy profile, with 36.8 ± 2.0% in *A. americana* and 33.3 ± 0.7% in *Antillogorgia* sp. 1 and correspondingly more area in the lower bands; *P. homomalla* (39.9 ± 1.9%) and *P. tenuis* (39.9 ± 1.4%) fell between these extremes (Online Fig. 2).

The profiles of the area occupied by each colony were presented as a function of height above the substratum to visualize space use. Relative band areas (i.e., the percentage of total colony area in each band) visualize the shape of the colony independent of height. Absolute area profiles depict the colonies’ use of space relative to distance above the substrate and followed a general pattern of increasing band area with increasing height above the substratum. In all species the greatest band area was below the topmost band, reflecting the tapered form of the colony (Fig. 4A). The relative colony profiles for any one species were broadly similar across sizes (Fig. 4B).

**Fig. 4.**
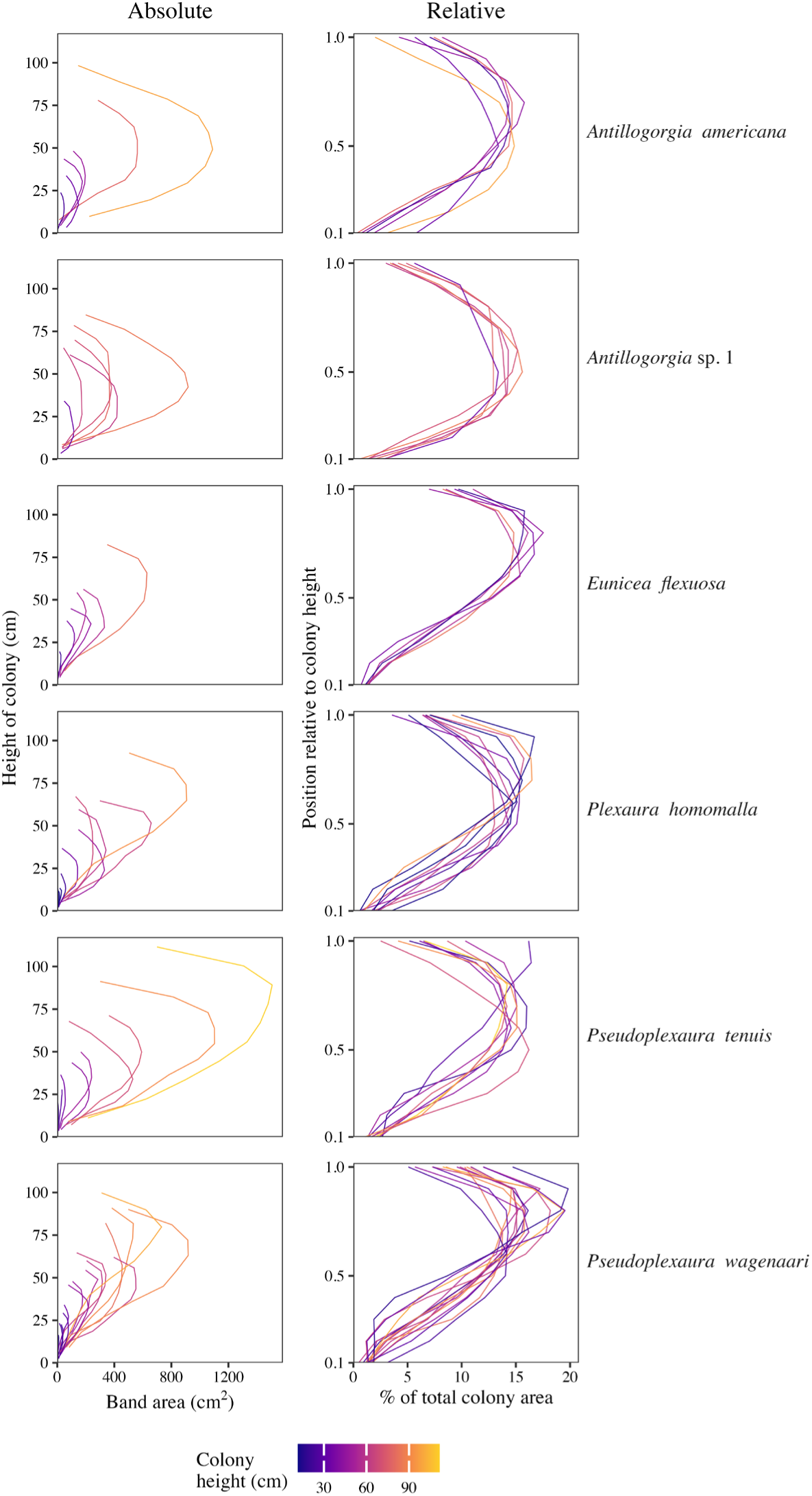
Absolute colony profiles to the left and relative colony profiles to the right for all six species.

Colony height had no detectable effect on relative colony shape. PC1 scores for relative band areas were unrelated to height (*R*^2^ = 0.015, *F*_1,50_ = 0.78, *p* = 0.38; slope = −0.012 PC1 units cm−1, 95% CI −0.041 to 0.016), as were PC2 scores (*R*^2^ = 0.013, *F*_1,50_ = 0.65, *p* = 0.42) (Fig. 5), and PERMANOVA on relative band areas returned no multivariate effect of height (*R*^2^ = 0.024, pseudo-*F*_1,50_ = 1.21, *p* = 0.27). With species in the model, height explained a further 1.2% of variation in relative morphology (pseudo-*F*1,45 = 0.93, *p* = 0.35), against 41.1% for species (pseudo-*F*_5,45_ = 6.54, *p* = 0.0001). Height × species interactions were not significant on either PC1 (*F*_5,40_ = 0.43, *p* = 0.82) or PC2 (*F*_5,40_ = 0.68, *p* = 0.64), and per-species slopes of PC1 on height ranged from −0.029 to 0.017 with every 95% confidence interval containing zero. Repeating the species and height PERMANOVAs on centered log-ratio transformed band areas gave the same result (species *R*^2^ = 0.33, *p* = 0.0002; height *R*^2^ = 0.031, *p* = 0.19), as did adding site to the model (species *R*^2^ = 0.42, *p* = 0.0001; site *R*^2^ = 0.032, *p* = 0.72).

**Fig. 5.**
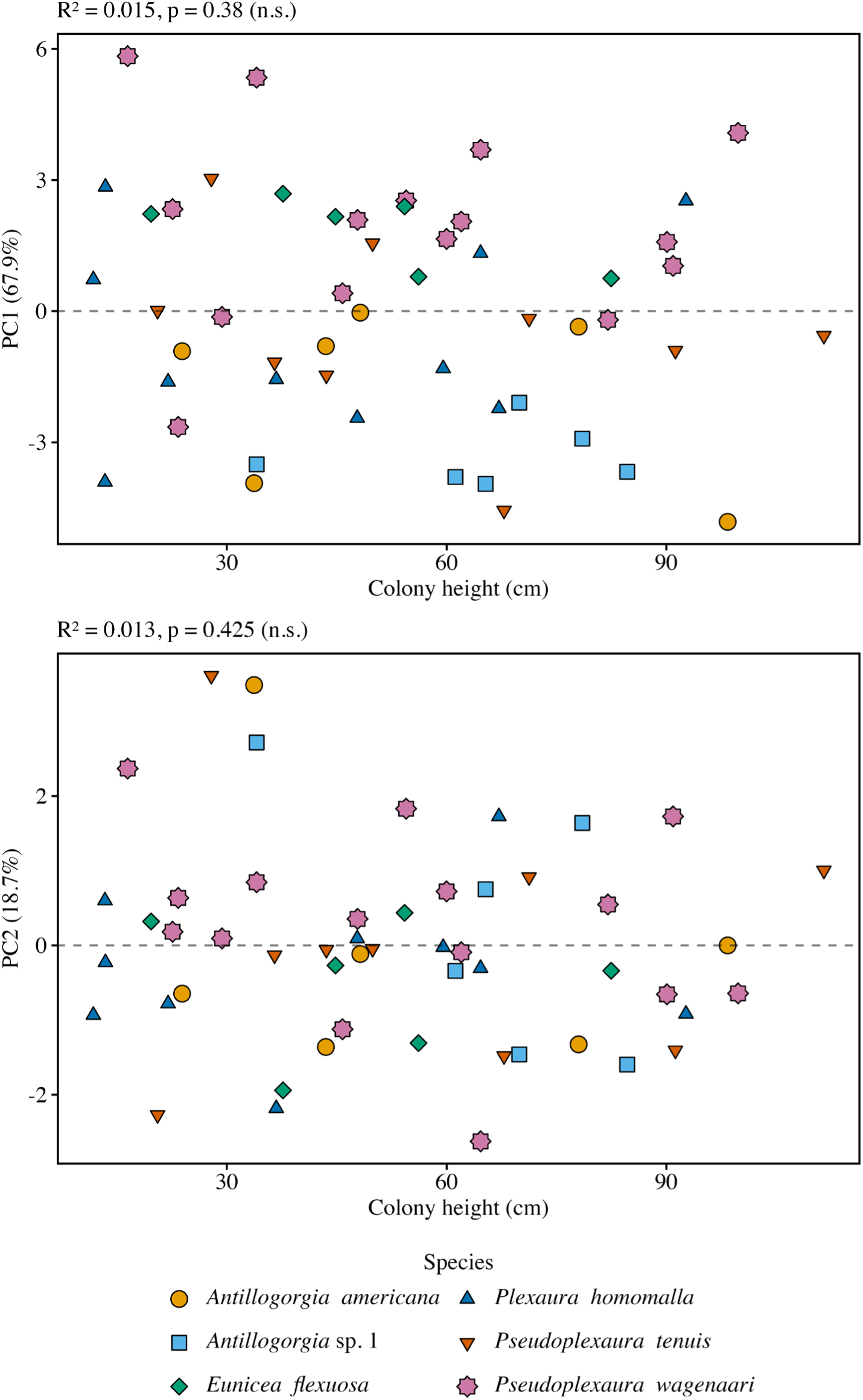
Relative band-area PC1 (top) and PC2 (bottom) vs. colony height. No fitted line is shown because height had no detectable effect on either axis.

Although overall form was unaffected by colony height, the proportion of a colony’s area occupied by branches increased with height (Fig. 6; slope = 0.0056 ± 0.0021 logit units cm−1, *F*1,46 = 6.91, *p* = 0.012), from a predicted 41% at 20 cm to 52% at 100 cm, and the increase did not differ among species (height × species interaction, *F*_5,36_ = 0.70, *p* = 0.63). Branch fraction also differed among sites (*F*_4,46_ = 4.52, *p* = 0.004). Colonies therefore grew without a detectable redistribution of area along the vertical axis, filling their outline more densely instead.

**Fig. 6.**
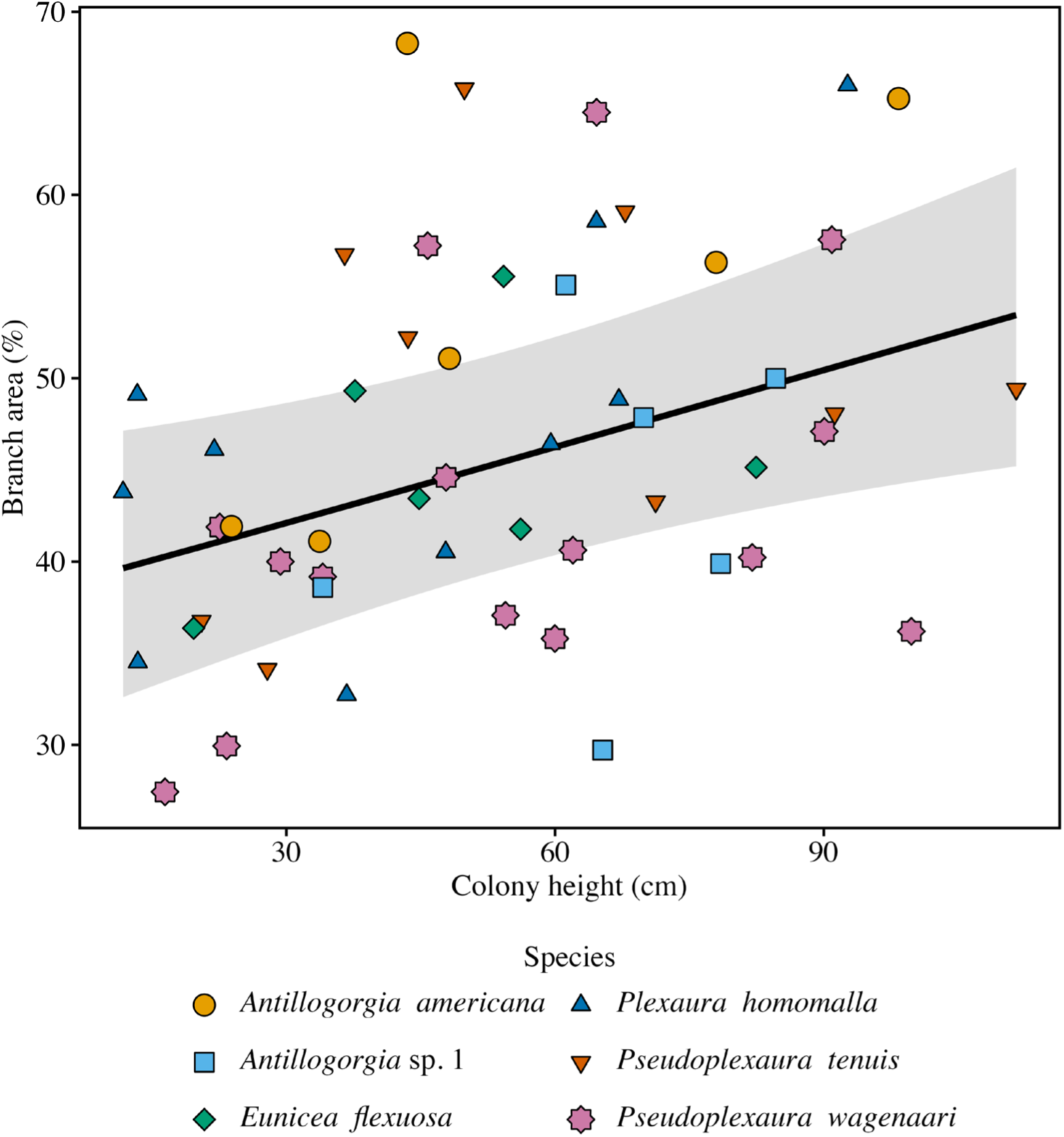
The percentage of a colony’s area occupied by branches modeled as a function of colony height for all six octocoral species. The fitted line is from a quasi-binomial generalized linear model that also included site as a factor.

## Discussion

As scleractinians decline in abundance on Caribbean reefs, the physical structure of the benthos is increasingly determined by the form of octocoral colonies, and that form is the product of colony growth. We asked two questions about colony growth in six sympatric species: whether the species differ in colony form independent of size, and whether relative form changes with colony height. The first was supported and the second was not. Species identity separated colonies along PC1 of the relative band-area PCA, the axis carrying most of the variation in relative form, while colony height did not shift scores on either axis. Together, these results indicate that the branching program of each species sets a distinctive distribution of area occupied by the colony along the vertical axis (see also Sánchez 2004), and that colonies retain that shape as they grow. This has implications for the structure of octocoral assemblages and for the broader question of how three-dimensional structure is generated in modular organisms.

Differences in colony shape among species fell almost entirely on the first axis of the PCA, which captured proportional differences in the distribution of area occupied along the colonies’ vertical axis. Pairwise PERMANOVA distinguished significant differences in eight of 15 species pairs after FDR correction. Those cases of significant difference included all of the pairings of the two *Antillogorgia* species with *E. flexuosa* and *P. wagenaari*. The differences among species on PC1 were consistent with the qualitative characterizations of the colonies’ architecture. *Antillogorgia* build dense, plumose canopies in which branches and associated branchlets are deployed throughout the height of the colony and, being quite flexible, often droop downward. Thus, the convex-hull outline contains substantial area in the lower and middle bands, and the projected profile of the branches is more uniformly distributed along the vertical axis than in the other species in the study (Fig. 4 and Online Fig. 2). *E. flexuosa* and *P. wagenaari* produce more open forms in which sparse upper branches fan outward from a primary branch that is comparatively bare close to the base, and those colonies carry a larger share of their projected area in the upper bands. Because this PCA was based on relative band areas, the differences among species observed on PC1 were not a byproduct of size. Small *A. americana and Antillogorgia* sp. 1 colonies appear to generate branches following the same rules that produce the plumose adult. Species pairs whose profiles were similar (e.g., P. *homomalla* and *P. tenuis*, or *E. flexuosa* and *P. wagenaari*) occupied adjacent positions on PC1. The PERMDISP test indicated that within-species variability was similar across the assemblage (*F*_5,46_ = 1.05, *p* = 0.40), so the failure to separate these pairs reflects genuine overlap in colony form rather than uneven dispersion in shape space. That pattern is consistent with the morphological gradient documented across the plexaurid genera, where branching density and colony shape vary continuously across closely related species (Brazeau and Lasker 1988; Kupfner Johnson and Hallock 2020).

The absence of a height effect on relative band areas is consistent with the self-organized branching model proposed for arborescent gorgonians (Sánchez et al. 2004b), in which axial extension and branching follow a structured sequence that sets a colony’s branching pattern as modules are added. (Mother branches produce branches as they elongate while daughter branches do not produce new branches.) Most notably, Sánchez et al. (2004b) show how colonies grow in a pattern that maintains the ratio of total to mother branches. That suggests a stasis of form. However, while the relative shape stays constant the colony’s profile does change. For instance, the absolute positioning of the widest band is higher in tall colonies than in small colonies (Fig. 4A). In work with *Antillogorgia elisabethae,* Lasker et al. (2003) found that extension of daughter branches declined with time and length of the branch, whereas extension of mother branches was greater than that of daughter branches. While the branch extension of mother branches declined in older and taller colonies, it continued to be greater than that of daughter branches. In combination those traits generate a colony form in which single plumes maintain their feather-like shape, and as new plumes develop, the acute angle of their growth relative to the branch they arose from means the height at which the colony is widest will increase as the colony grows. As the number and length of mother branches increase, the size of the polygon increases. Furthermore, the increasing number of mother branches produces an increasing number of daughter branches which fill in the space of the polygon, consistent with the increase in branch fraction with height (Fig. 6). The rules governing branching, the creation of mother branches and extension rates are a developmental program. The overall growth of the colony and the maintenance of its shape are consequences of those rules.

Our findings are also relevant to the structure of the octocoral forests that increasingly dominate Caribbean reefs (Lasker et al. 2026). Because architecture is species-specific, the vertical distribution of canopy area in a multispecies stand would depend on its mixture of plumose and whip-like colonies and not simply on the sizes of those colonies. As evident from Fig. 4A, the profiles of the colonies grow as height increases, and the space also fills in through branching within the colony outline (Fig. 6). Thus, the canopy effects of an octocoral forest might be partitioned into a species-composition component that governs the shape of the canopy and a second component, reflecting both colony size and site, that governs its density. This three-dimensional structure of the octocoral forest matters for the ecological role these forests are now playing on Caribbean reefs (Lasker et al. 2020). Near-bed seawater flow and turbulence in animal forests are sensitive to canopy density and vertical structure (Guizien and Ghisalberti 2017) and recruitment in the understory of octocoral assemblages is associated with the density of adult colonies on the reef (Privitera-Johnson et al. 2015). Decomposing canopy structure in this way is the same approach used to study how forest stands modify their environment as they mature (Nelson and Bramanti 2020), and it provides a vocabulary for treating octocoral forests as analogues of arborescent plant canopies. Applying PCA to the stacked horizontal band quantification of colony shape of these arborescent colonial animals also shows that the cross-sectional approach used in plant allometry (King 1990; Enquist 2002) can be extended to colonial cnidarians, and is in principle applicable to other modular taxa, including sponges, bryozoans, hydroids, and large macroalgae, in which colony or thallus form is generated by iterative module addition (Harper 1977; Kaandorp and Kübler 2001). The contrast with the cross-sectional allometric framework used in plant ecology is instructive. In trees, increased height is accompanied by predictable changes in the distribution of crown area and in the proportion of the standing structure that is photosynthetically active (King 1990; Enquist 2002). The octocorals here showed no such redistribution, and added branches within a profile whose proportions did not detectably change with size.

Multiple lines of work would extend the present analysis and add to our understanding of the ecological role octocorals play on modern reefs. Three-dimensional photogrammetry would provide volumetric profiles, creating a more complete characterization of form by adding a third dimension. Longitudinal tracking of tagged colonies would test whether the size-independence of relative form inferred here across colonies also holds within a colony followed through time. Whether the vertical distribution of branches on recovering colonies converges on the species’ PC1 position after partial mortality and branch loss is also unresolved (Castanaro and Lasker 2003; Sánchez and Lasker 2004a; Anderson et al. 2022). Colony form will also respond to variation in habitat and depth among sites (Kim et al. 2004; Prada et al. 2008), so partitioning genotype-by-environment effects on architecture is a natural extension. Expanding the comparison to taxa with a broader array of species and colony shapes, such as *Gorgonia* spp. and other *Antillogorgia* spp., would test whether the independence of species identity and growth is a general property of Caribbean octocorals and whether differences among species also have a phylogenetic signal. It is also worth noting that the 95% confidence interval on the pooled slope of PC1 on height admits a shift of up to 4.1 PC1 units across the 99.7 cm range of colony height sampled here, against a spread of 5.3 units among species means, so a size effect smaller than the differences among species would have gone undetected. Sampling a wider size range within each species would test it directly. Finally, linking PC1 position to measures of functional performance under differing flow and feeding regimes (Sarmiento et al. 2026) would help identify the ecological significance of the species’ respective modular growth programs.

## Data Availability

The morphometric dataset and the R analysis code supporting these findings are available from the authors and will be deposited in a public repository on acceptance.

## Acknowledgements

We thank J. Benz and E.A. Anderson for assistance in the field and the Virgin Islands Environmental Resource Center for facilitating the work, as well as P.J. Edmunds for comments on the research and manuscript. This research was funded by the National Science Foundation (grant OCE 1756381) and conducted under research permits from the U.S. National Park Service (VIIS-2021-SCI-0008).

## Notes

### Competing Interest Statement

The authors have declared no competing interest.

