## Supplementary material for "Species identity rather than colony size partitions shape variation in six Caribbean octocorals": Online Material

**Online Supplemental Material**

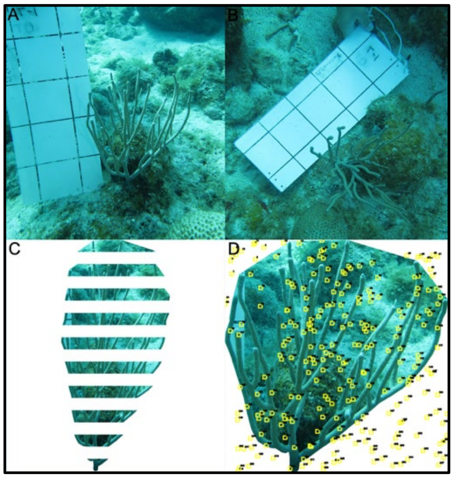


**Online Fig. 1** Steps involved in the analysis of images and collection of morphometric data for a colony of *Pseudoplexaura wagenaari*. A, Initial image with a 10 × 10 cm scaling grid; B, Colony polygon; C, Colony band; D, CoralNet point counts

**Online Fig. 2** Average relative colony profiles of all 6 octocorals used in the study. *Antillogorgia americana* and *Antillogorgia sp. 1* shared a more bottom-heavy profile, *Eunicea flexuosa* and *Pseudoplexaura wagenaari* were top-heavy, and *Plexaura homomalla* and *Pseudoplexaura tenuis* were intermediate.
